# Modern wheat breeding selection synergistically improves above- and below-ground traits

**DOI:** 10.1101/2023.10.27.564104

**Authors:** Peng Zhao, Zihui Liu, Xue Shi, Wenyang Hou, Mingzhu Cheng, Yuxiu Liu, James Simmonds, Wanquan Ji, Cristobal Uauy, Shengbao Xu, Xiaoming Wang

## Abstract

The root system, as a fundamental organ for uptaking water and nutrients and interacting with the local environmental conditions, has been postulated to be the foundation for a second Green Revolution. However, the status of the root system during modern wheat breeding remains to be elucidated. Here, by analyzing the phenotypes of 406 wheat accessions on a large scale, we found the root systems of modern cultivars were synchronisely changed along with the above-ground traits. Furthermore, the genomic blocks with phenotypic effects on both above- and below-ground traits were observed to be enriched in the selection sweeps, highlighting that modern wheat breeding contributed to the synchronised changes. More importantly, the haplotypes selected by breeders within the selection sweeps synergistically improved both the above- and below-ground traits, suggesting that modern wheat breeding has improved the root system indirectly, which may contribute to the higher grain yields of modern wheat cultivars. Our results demonstrated that modern wheat breeding synergistically improved the above- and below-ground traits.

---

The root system is a fundamental organ for the uptake of water and nutrients from the soil and defines the interaction of the plant with the local environment (Calleja-Cabrera et al., 2020). As such, it plays a vital role in meeting the increasing global food demand and has been proposed as the critical target for the second Green Revolution (Herder et al., 2010). Bread wheat (*Triticum aestivum* L.) is the most widely grown crop and provides one-fifth of the total calories of the world’s population (FAO, 2023). Over the past century, wheat breeding has remarkably increased yield by selecting for above-ground traits (Tadesse et al., 2019; Voss-Fels et al., 2019). However, the effects on the root system remain largely unknown.

Previously, we found that modern wheat cultivars (MC) have larger root systems than landraces (LA), including root surface, root volume, fresh root weight and root diameter, by phenotyping and transcriptome-sequencing of a panel of 406 worldwide accessions (Wang et al., 2022). These results were also consistently observed when the confounding effects of kernel weight were considered (Supplemental Figure 1 and Supplemental Data 1), suggesting modern wheat breeding also changed the below-ground traits, albeit indirectly. We then phenotyped the above-ground traits in seven environments. As expected, all the investigated traits were significantly changed in the MC group compared with the LA group, except for spikelet number (Supplemental Figure 1). More importantly, the variation of above-ground traits was significantly correlated with that of the below-ground traits among the population, especially for plant architecture and kernel-related traits, which were the strongest targets of breeding selection (Figure 1A and Supplemental Figure 2A). Interestingly, these correlations emerged or were more significant in the MC group than in the LA group (Figure 1B and Supplemental Figure 2B). These results showed that wheat breeding synchronously changed the above- and below-ground traits.

**Figure 1.**
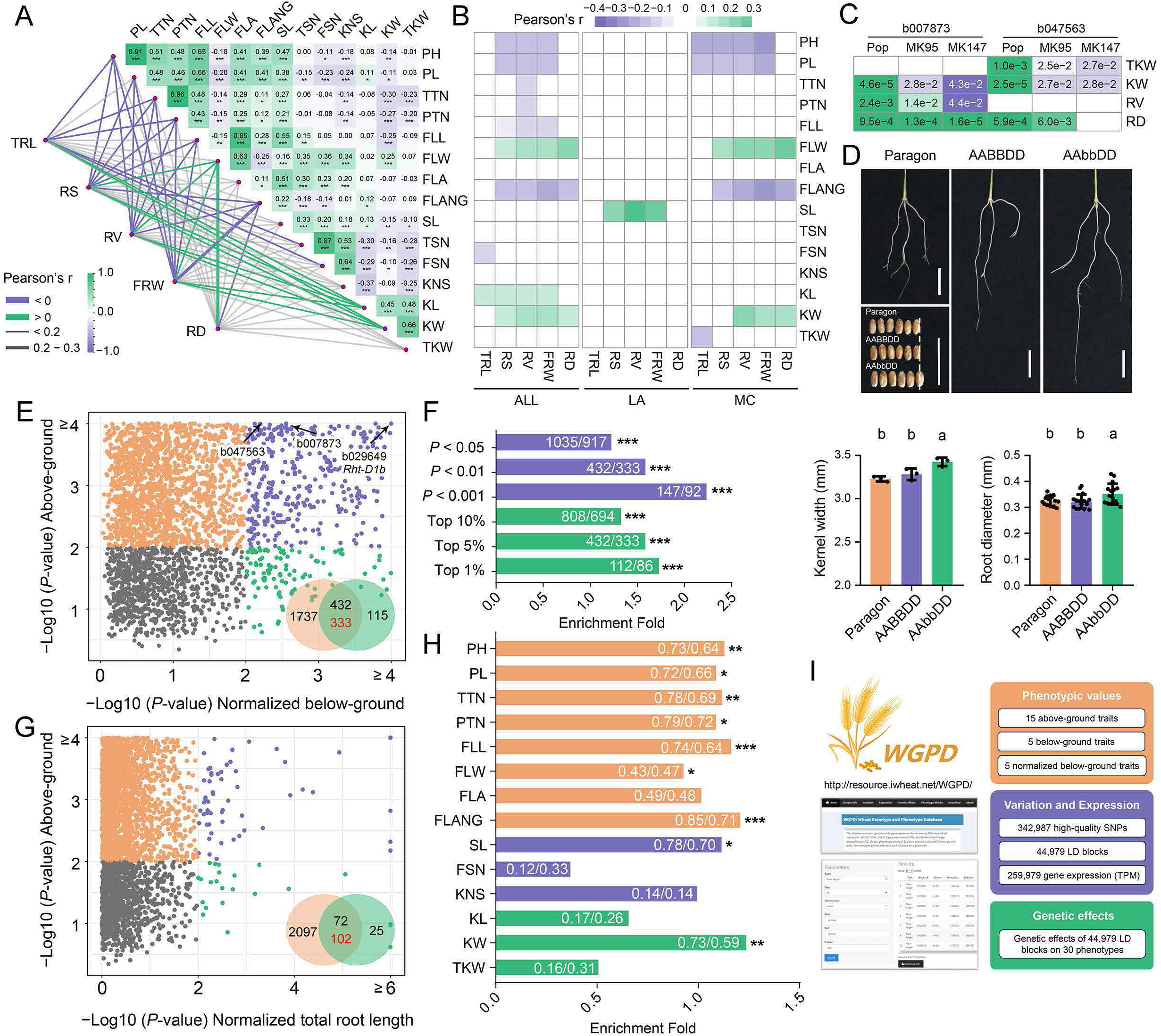
Synergistic improvements between above- and below-ground traits during wheat breeding selection. (A) Correlation between above- and below-ground traits in wheat. The phenotype values of below-ground traits were normalized with the general linear model to exclude the effects of the kernel weight. The same results were also observed with the original phenotype values (Supplemental Data 1 and Supplemental Figure 2A). The left section indicates the correlation coefficients and the root traits (TRL: total root length; RS: root surface; RV: root volume; FRW: fresh root weight; RD: root diameter). The right section indicates the correlation among above-ground traits. The short name of the traits represents PH (plant height), PL (peduncle length), TTN (total tiller number), PTN (productive tiller number), FLL (flag leaf length), FLW (flag leaf width), FLA (flag leaf area), FLANG (flag leaf angle), SL (spike length), TSN (total spikelet number), FSN (fertile spikelet number), KNS (kernel number per spike), KL (kernel length), KW (kernel width), TKW (thousand kernel weight). The potential effects of population structure on the correlations were corrected with the general linear model. (B) The correlation between above- and normalized below-ground traits in the landraces (LA) group, modern cultivars (MC) group and all accessions. The potential effects of population structure on the correlations were corrected with the general linear model. (C) Genetic effect validations of the bi-effect blocks b007873 and b047563. The relative phenotype values of the individuals carrying alternative haplotypes in three populations were compared, and the number on the square represents the significance *P*-value. Labels indicate the whole association population (Pop), RIL population derived from crossing between MK95 and CS (MK95), and RIL population derived from crossing between MK147 and CS (MK147). Additional information is in Supplemental Figure 3 and 4. (D) The root and kernel phenotype of *TaGW2* gene mutants. Wild type (Paragon and AABBDD) and single mutants (AAbbDD). Data are means ± SD; letters on top indicate statistical differences (*P*-value < 0.05) based on the least significant difference (LSD) test. Root phenotypes were evaluated at 14 days after germination. Bar = 2 cm. Additional information is in Supplemental Figure 5. (E) The enrichment of bi-effect blocks in the selection sweeps. Each point represents a selection block. The *x*- and *y*-axis indicate the *P-*value of each selection block in the genetic effect estimations. The arrows indicate the blocks used to validate the genetic effects. In the Venn diagrams, the orange and green sections represent the selection blocks which had genetic effects on the above- and below-ground traits, respectively. The overlap represents the bi-effect blocks. The numbers in black are the observed values, while the maximum of the expected values in the 1,000 permutation tests are in red. (F) The fold-enrichment of bi-effect blocks in the selection sweeps increased with more stringent thresholds, from *P-*value 0.05 to 0.001 in the genetic effect estimation and from top 10% to 1% in the selection sweep detections. For the number pair on each bar, the left number is the observed value, and the right number is the maximum of the expected values in the 1,000 permutation tests. (G) The bi-effect blocks between above-ground traits and total root length were not enriched in the selection blocks. (H) Breeding selection synergistically improves the above- and below-ground traits. For the number pair on each bar, the left number is the ratio of bi-effect selection blocks with synergistic improvement effects to the bi-effect selection blocks with genetic effects on relative above-ground traits; the right number is the ratio of genome-wide bi-effect blocks with synergistic improvement effects to the genome-wide bi-effect blocks with genetic effects on relative above-ground traits. Pearson’s Chi-squared test *P-*value of < 0.05, < 0.01 and < 0.001, are indicated by asterisks (*, ** and ***), respectively. (I) Overview of the Wheat Genotype and Phenotype Database.

To explore the mechanisms underlying the synchronised changes, we estimated the genetic effects of genome-wide linkage disequilibrium (LD) blocks on each trait by analyzing whether phenotypes are significantly different among the haplotypes in a given LD block with the linear mixed model (Supplemental Data 2). A total of 3,614 blocks (out of 44,979 blocks) were identified as bi-effect blocks, which had genetic effects on both above- and below-ground traits, providing genetic clues for the synchronized changes. To experimentally validate the dual roles of the bi-effect blocks, firstly, two blocks (b007873 and b047563) were randomly selected, and their genetic effects were validated with two F_5:6_ segregation populations (Figure 1C, Supplemental Figure 3 and 4). Then, the block b041231, containing the well-known *TaGW2-B1* (*Grain Width 2*) (Wang et al., 2018), was identified to have genetic effects on both kernel width (*P*-value = 5.88e-3) and root diameter (*P*-value = 1.34e-3). Consistently, the mutants of *TaGW2-B1* showed significantly increased kernel width and root diameter compared with the wildtypes (Figure 1D and Supplemental Figure 5). Lastly, the block b029649 showing the most significance level contains the Green Revolution gene *Rht-D1 (Reduced Height D1)*, where its *Rht-D1b* allele was demonstrated to have genetic effects on both above- and below-ground traits in our previous studies (Liu et al., 2022; Wang et al., 2022). These results provide strong evidence for the reliability of the bi-effect blocks.

We next identified 2,461 selective sweeps with the top 5% of *F*_st_, π ratio, and XP-CLR (Supplemental Data 3) covering 3,389 LD blocks (named as selection blocks hereafter) and including several well-known selected genes (Supplemental Data 4). The percentage of bi-effect blocks in the selection sweeps (432 vs. 3,389, 12.75%) was higher than that in the genome-wide background (3,614 vs. 44,979, 8.03%), demonstrating that the bi-effect blocks were enriched in the selection sweeps (Pearson’s Chi-squared test *P-*value = 1.29e-25) (Figure 1E). This suggests that modern breeding contributed to the synchronized changes, which was consistent with the higher correlations between above- and below-ground traits in the MC group.

To exclude the possibility of this enrichment arising by chance, we performed 1,000 permutation tests and recorded the number of bi-effect blocks in each test. The numbers ranged from 223 to 333, with an average of 272, which was significantly smaller than the observed value (n□= 432, *P-*value < 0.001) (Supplemental Figure 6). Meanwhile, the enrichment folds increased from 1.23 to 1.59 and 2.23 by increasing the *P-*value threshold in the genetic effects estimation from 0.05 to 0.01 and 0.001, respectively. They also increased from 1.33 to 1.59 and 1.73 by increasing the threshold in the selection sweep detection from the top 10% to 5% and 1%, respectively, despite a rapid drop in the number of bi-effect blocks (Figure 1F and Supplemental Figure 7). In contrast, the bi-effect blocks between above-ground traits and the total root length that had no difference between LA and MC groups were not enriched in the selection blocks (Figure 1G and Supplemental Figure 8). Together, these results provide strong evidence that wheat breeding acted on the bi-effect blocks, thereby contributing to the synchronized changes.

The synchronized changes driven by breeding selection prompted us to ask whether the selected bi-effect blocks synergistically improved above- and below-ground traits toward directions of the MC group. To address this question, we considered the haplotypes with increased frequency in the MC group in each of the bi-effect selection blocks as selected haplotypes. We then queried if the selected haplotypes improved both the above- and below-ground traits (synergistic effects) compared with the genetic effects of the remaining, non-selected haplotypes in each block. A total of 392 out of 432 (90.74%) bi-effect selection blocks, were identified to have synergistic improvement effects (Supplemental Data 5); for example, the selected haplotypes in 198 bi-effect selection blocks simultaneously reduced plant height and increased root systems (root surface, volume, fresh weight, or diameter). Moreover, the ratio of bi-effect selection blocks with synergistic improvement effects was larger than the relative ratio under a genome-wide background, except for the analysis from flag leaf width, fertile spikelet number, kernel length, and thousand kernel weight (Figure 1H). These results prove that wheat breeding synergistically improved above- and below-ground traits by selecting the haplotypes with synergistic effects.

To understand the underlying mechanisms of the synergistic effects, we identified genome-wide expression QTLs (eQTLs) and found 471 eQTLs located within 281 bi-effect selection blocks regulating the expression of 1,862 genes (Supplemental Data 6). A transcriptome-wide association study (TWAS) revealed that the expression levels of 276 genes, regulated by 189 eQTLs (40.13%) from 151 bi-effect selection blocks (53.74%), were significantly associated with below-ground trait variations (FDR < 0.01; Supplemental Data 7), providing a connection between bi-effect selection blocks and the below-ground trait changes. Among these 276 genes, we identified wheat homologs for genes known to be essential in root development of model plant species, including *OsGCN5, AtMNS3, AtPERK8* and *AtPGP4* (Supplemental Figure 9). Similarly, the ratio of bi-effect selection blocks regulating significantly associated genes in TWAS was higher than the expected value under the genome-wide background (53.74% vs 39.86%).

In summary, we demonstrated that modern wheat breeding synergistically improved the above- and below-ground traits through the selection of haplotypes with synergistic effects in the bi-effect selection blocks. We established the Wheat Genotype and Phenotype Database (WGPD, http://resource.iwheat.net/WGPD/), a free, web-accessible, and user-friendly platform. In the WGPD, users can freely download the genotype, gene expression, and phenotype data used in this analysis and search for the genetic effects of a given LD block and the related LD blocks for a given trait (Figure 1I).

## Author contributions

X.W., S.X., and C.U. conceptualized and designed the study. P.Z. and X.S. performed the data analysis. Z.L. measured the phenotypes of the above-ground traits. J.S. created the mutants of *TaGW2*. W.H. and M.C. collected the genotype and phenotype data of the two F_5:6_ segregation populations and conducted phenotyping of the *TaGW2* mutants. Y.L. collected the well-known selected genes. X.W., S.X., and C.U. wrote the manuscript. All authors discussed the results and approved the final version of the manuscript.

## Acknowledgments

We thank the Prof. Shifeng Cheng (Agricultural Genomics Institute at Shenzhen, Chinese Academy of Agricultural Sciences) for careful proofreading and comments on the manuscript. And thank the computing platform support of High-Performance Computing of NWAFU. No conflict of interest is declared.

