## Supplemental Information for "Modern wheat breeding selection synergistically improves above- and below-ground traits"

**Materials and Methods**

**Plant materials and** **phenotypic data**

The 87 landraces and 190 modern cultivars used in this analysis (Supplemental Data 1), the phenotyping data of root-related traits and the genotypes of each accession were from our previously reported association population composed of 406 bread wheat accessions (Wang et al., 2022). Briefly, the root development traits 14 days after germination, including total root length (TRL), root surface (RS), root volume (RV), fresh root weight (FRW) and root diameter (RD), were measured with six independent replications and at least six plants with consistent development status were measured in each replication. At the same time, one plant with consistent development status as that used for root phenotyping was selected from each replication, and its root sample was frozen with liquid N_2_. The root traits were normalized with a general linear model and the “lm” function in R (v 3.6.1) to exclude the effects of kernel weight on root-related traits. Then, six independent samples from six replications were equally bulked for pair-end RNA-seq with an insert size of ∼250 bp on the Illumina HiSeq X Ten platform. The clean reads were mapped to the reference genome of Chinese Spring (IWGSC RefSeq v1.0) (IWGSC, 2018) and the annotated transcripts (IWGSC RefSeq v1.1). The mapping results were passed to the pipeline developed and refined to improve the accuracy of SNP detection and gene expression quantitation (https://github.com/biozhp/Population_RNA-seq) (Wang et al., 2022), generating 1,232,311 high-quality SNPs and the expression levels of 107,662 genes among the population. The above-ground traits were measured in seven environments at Yangling (34.28'N, 108.07'E, altitude 517 m) and Chongzhou (30.63'N, 103.67'E, altitude 1300 m) in China in 2018-2021 (Liu et al., 2022). All the accessions were planted in one-row plots (1 m in length and a row spacing of 20 cm) in an incompletely randomized block design with three replicates in each environment, and nine plants were measured in each replication. The best linear unbiased estimate (BLUE) values under seven environments were obtained with a linear mixed model in which genotypes and replicates were added as the fixed effect and random effects, respectively.

**Genomic LD block construction and genetic effect estimation**

The genomic LD block construction used methods reported in (Voss-Fels et al., 2019). SNPs with minor allele frequency (MAF) > 0.05 were assigned to LD blocks based on pairwise *r*^2^ > 0.7. Both the first step of pairwise *r*^2^ value calculating and the initial LD block constructions and the second step of the LD block adjustments were performed in the R package SelectionTools (v22.1) with the parameter (ld.threshold = 0.7, ld.criterion = "flanking", tolerance = 1). The SNPs not in LD with any other SNPs were assigned to individual LD blocks.

The haplotypes within each LD block were defined with SNP combinations. Only haplotypes present in more than 30 accessions were selected for the genetic effect estimation. The LD block would be regarded as having genetic effects on a given trait when the phenotyping values of accessions carrying different haplotypes within this LD block were significantly different. The significance was calculated using a linear mixed model with the lmerTest (v3.1.3) package in R (v 4.2.2), in which the population structure and familial relatedness were included as fixed and random effects, respectively, to exclude their interferences with the following equation:

$$y =\mu+ Xa + P\beta+ K\gamma+ e$$

In this equation, *y* represents the phenotype values of the investigated accessions, *μ* represents the overall mean, and *Χ* represents the haplotypes within the investigated LD block. The *P* and *K* parameters represent the PCA and the relative kinship matrix generated with GAPIT (v3.1), respectively. The top three principal components were used to build the *P* matrix for population-structure correction. The *K* matrix was used to correct the potential familial relatedness. *Xα* and *Pβ* represent fixed effects, and *Κγ* represents random effects. The significance (*P-*value) of *Xα* was extracted using the "anova" function of the R (v 4.2.2). The LD blocks which had genetic effects on at least one above-ground trait and at least one below-ground trait was defined as bi-effect blocks.

**Pearson’s correlation analysis**

The phenotypic values are regressed on population structure using a linear model with the “lm” function in R (v 4.2.2), aiming to exclude the potential effects of population structure on correlation analysis. Then, the adjusted phenotypic values (regression coefficients + residuals) were passed to Pearson’s correlation test using the “cor.test” function in (v 4.2.2).

**Identification of selective sweeps**

SNPs with MAF > 0.05 were used to calculate the nucleotide diversity (π) and genetic differentiation (*F*_st_) between MC and LA groups using VCFtools (v0.1.16) with 200-kb sliding windows and 100-kb steps. The XP-CLR score between MC and LA groups were calculated using XP-CLR with 200-kb sliding windows and 100-kb steps. The windows with the top 5% of π ratio (π_LA_/π_MC_), *F*_st_ and XP-CLR scores were considered selective sweeps during wheat breeding. The different thresholds regarding the top percentages (top 1% and top 10%) were also used for the relative analysis, and consistent results were observed among them (Figure 1F and Supplemental Figure 7B).

**Genetic effect validation of the bi-effective blocks with segregation populations**

Two F_5:6_ recombinant inbred line (RIL) populations composed of 121 lines and 103 lines were derived from crossing between Chinese Spring (CS) and MK95 and CS and MK147, respectively. The populations were developed by single seed descent and advanced to the F_5:6_ generation. The two F_5:6_ RIL populations were planted in one-row plots (1 m in length and a row spacing of 20 cm) with three replications in 2021-2022, and the main stem spikes from the three plants in the middle of each line were harvested for the phenotyping of kernel-related traits. Then, the harvested seeds were germinated, and their root-related traits were measured with the same procedure as that used for the association population phenotyping. For genotyping, one Kompetitive Allele Specific PCR (KASP) marker was developed for each of the two bi-effect blocks of b007873 and b047563, using PolyMarker (http://www.polymarker.info/) (Ramirez-Gonzalez et al., 2015) and used for haplotype assignments of the individuals among the two RIL populations (Supplemental Data 8). Lastly, for a given trait, the phenotype values of the individuals carrying alternative haplotypes were compared with the Student’s *t*-test. Only the homozygous lines were selected for the phenotype comparisons.

**Phenotypic evaluation of *TaGW2* gene mutants**

The loss-of-function mutants of the *TaGW2*, including Wild type (Paragon and AABBDD), single mutants (aaBBDD, AAbbDD, and AABBdd) and triple mutant (aabbdd), were derived from BC_4_ near-isogenic lines produced by crossing and back-crossing of the TILLING mutant lines (Kronos2335 for *TaGW2-A1*, Kronos0341 for *TaGW2-B1*, Cadenza1441 for *TaGW2-D1*) to the cultivar Paragon based on our previous study (Wang et al., 2018). The germinated seeds were cultured with damp filter papers for three days and then moved to containers (40×30×12 cm) filled with 1/2 Hoagland solution with punched lightproof plastic covers. The Hoagland solution was changed weekly. Root phenotypes were evaluated at 14 days after germination.

**Identification of genome-wide expression QTLs (eQTLs)**

The genes with expression levels more than 0.5 (TPM > 0.5) in more than 95% of accessions were selected and used for eQTL identification. Firstly, the expression values among the population for each gene were transformed to a normal distribution with the “qqnorm” function in R (v 3.6.1). Then, the hidden and confounding factors contributing to the expression variability were estimated with Bayesian factor analysis implemented in the PEER (v1.0). Then, the linear mixed models implemented in the GEMMA (v0.98.3) and MatrixEQTL (v2.3) packages that consider the population structure, genetic relatedness and the estimated confounding factors were used for association analysis between the SNPs with MAF > 0.05 (genotypes) and the normalized expression profile of a given gene (expression trait). The significantly associated SNPs (*P-*value < 6.367e-8, determined by 0.01/Total SNP number) generated by the GEMMA and the MatrixEQTL were intersected to improve reliability and accuracy.

To deal with the association of multiple SNPs with one expression trait and define the eQTL intervals, a revised two-step method (Fu et al., 2013) was employed as follows: (1) all of the associated SNPs were grouped into one cluster if the distance between two consecutive SNPs was less than 10 kb, and the clusters with at least three significantly associated SNPs were considered as candidate eQTLs represented by their lead SNP; (2) A candidate eQTL in LD (*r*^2^ > 0.1) with other more significant candidate eQTLs for the same expression trait were regarded as associations introduced by the LD structure and were removed.

**Transcriptome-wide association study (TWAS)**

The TWAS, aiming to calculate the association between the gene expression variation and the phenotype variation among the population, is usually performed with the software developed for genome-wide association analysis (GWAS) in plants by transforming the continuous variable (gene expression levels) into a discrete variable ("0" and "1"). However, this transformation results in losing part of the gene expression information. Here, we used the mixed linear model, the fundamental model in GWAS, to exploit the continuous gene expression variations and exclude the inference of population structure and family relatedness with the lmerTest (v3.1.3) package in R (v 4.2.2) as follows:

$$y =\mu+ Xa + P\beta+ K\gamma+ e$$

In this equation, *y* represents the values of the investigated phenotype, *μ* represents the overall mean, *Χ* is a continuous variable and represents the gene expression values. The *P* and *K* are the PCA and relative kinship matrix generated with GAPIT (v3.1) using the parameters “PCA.total=3”. The top three principal components were used to build the *P* matrix for population-structure correction. The *K* matrix was used to correct the potential familial relatedness. *Xα* and *Pβ* were used as fixed effects, and *Κγ* was used as random effects, respectively. The genes with false discovery rate (FDR) adjusted *P-*value < 0.01 were considered candidates significantly associated with the investigated phenotypes. Only the genes with expression levels higher than 0.5 (TPM > 0.5) in more than 95% of accessions were used for the TWAS analysis.

**Data availability**

All the accession information, genomics data and phenotypic data can be downloaded from our database (http://resource.iwheat.net/WGPD/). All the code used in the study is available on GitHub (https://github.com/biozhp/selection_improve).

**Supplemental Figures**

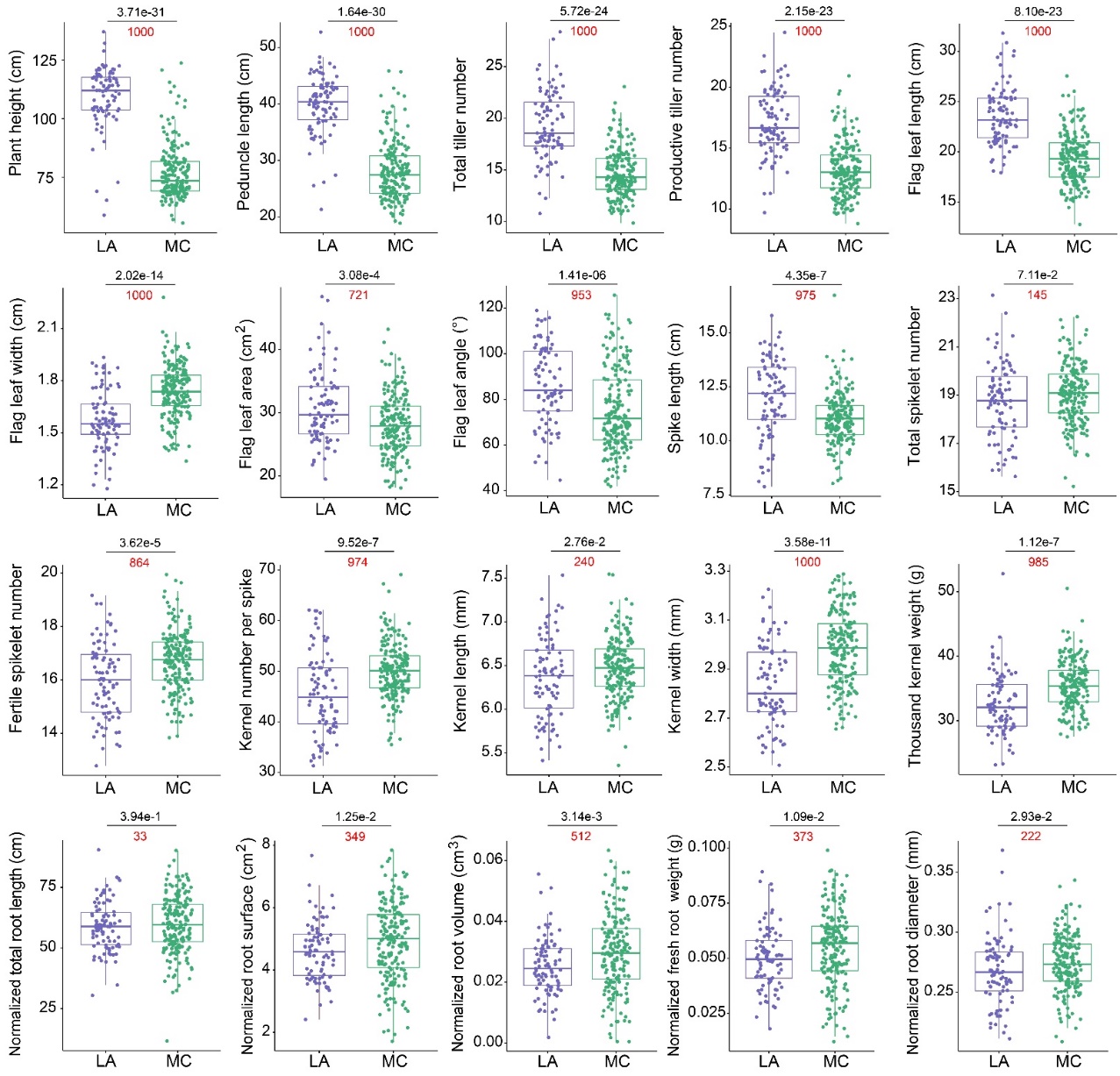

**Supplemental Figure 1. Comparisons of the above- and below-ground traits between landrace (LA) and modern cultivar (MC) groups.** The black numbers above each graph indicate the *P-*value from the Wilcoxon rank sum and signed rank tests. The red numbers show the observed trials with significant differences (*P-*value < 0.05) among 1,000 permutation tests. The permutation tests were performed by randomly sampling 52 accessions (60% of the LA accessions) from each LA and MC group and calculating the significance with Wilcoxon rank sum and signed rank tests. The phenotypic values of below-ground traits were normalized with the general linear model to exclude the effects of the kernel weight.

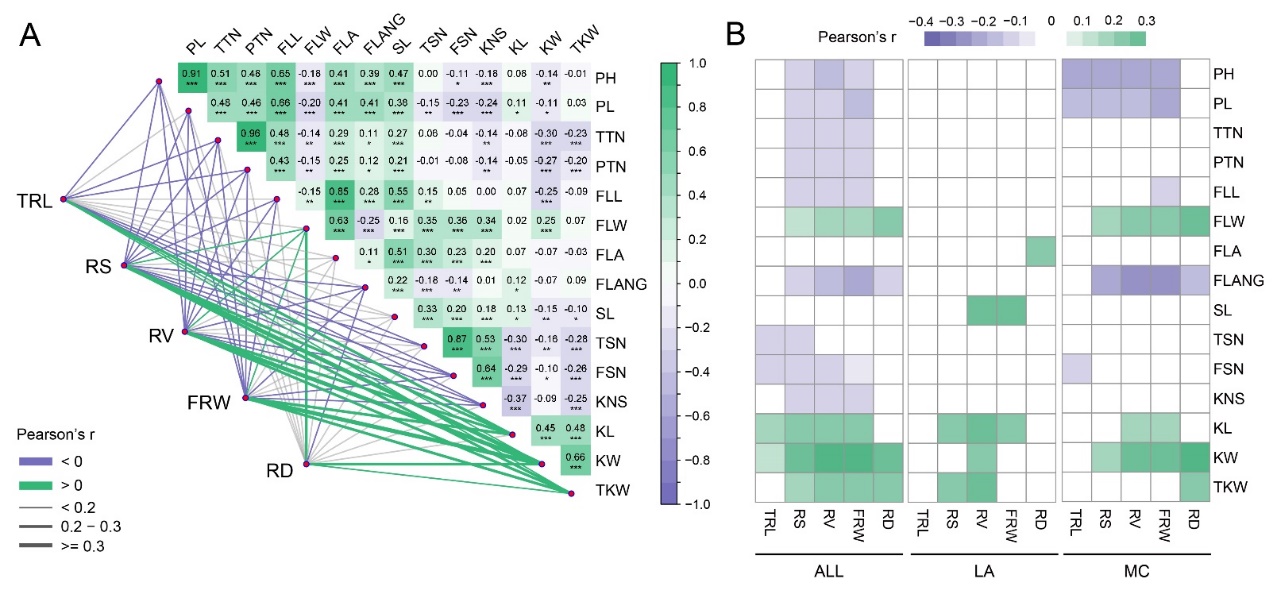

**Supplemental Figure 2. The correlation between above- and below-ground traits (original values).** (A) The original phenotype values of below-ground traits (uncorrected for kernel weight) were also significantly correlated with those of above-ground traits. The left part indicates the correlation coefficients between the above- and below-ground traits. The right part indicates the correlation coefficients among above-ground traits. The short name of the traits represents PH (plant height), PL (peduncle length), TTN (total tiller number), PTN (productive tiller number), FLL (flag leaf length), FLW (flag leaf width), FLA (flag leaf area), FLANG (flag leaf angle), SL (spike length), TSN (total spikelet number), FSN (fertile spikelet number), KNS (kernel number per spike), KL (kernel length), KW (kernel width), and TKW (thousand kernel weight). (B) The correlation between above- and below-ground traits (original values) in the landraces (LA) group, modern cultivars (MC) group and all lines. The potential effects of population structure on the correlations were corrected with the general linear model.

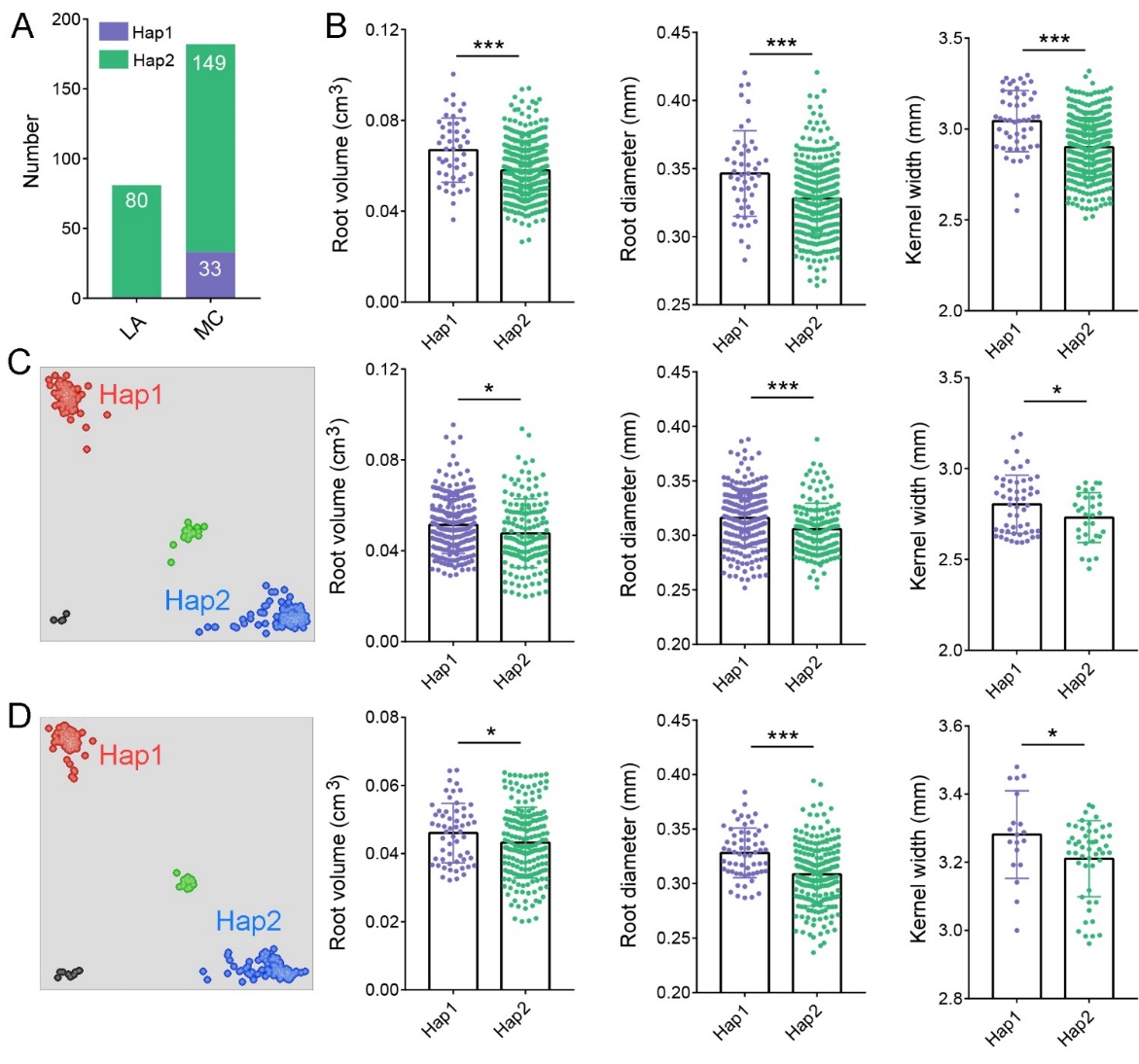

**Supplemental Figure 3. Validation of the genetic effects of bi-effect block b007873 on below-ground traits and kernel width.** (A) Frequency of Hap1 and Hap2 in LA and MC groups. (B) The phenotypic value distributions and significance values (Student's *t*-tests) between genotypes carrying Hap1 and Hap2 of block b007873 in the natural association population (top panels); F_5:6_ RIL populations derived from crosses between MK95 (Hap 1) and Chinese Spring (Hap2) (middle panels) and between MK147 (Hap 1) and Chinese Spring (Hap2) (lower panels). Asterisks indicate significance: * < 0.05; *** < 0.001. (C-D) The KASP assays to distinguish Hap1 and Hap2 genotypes in F_5:6_ RIL populations derived from crosses between (C) MK95 (Hap 1) and Chinese Spring (Hap2) and (D) between MK147 (Hap 1) and Chinese Spring (Hap2). For each F_5:6_ RIL population, five to eight plants of each line were genotyped, and the homozygous plants were phenotyped for the root-related traits. The homozygous lines were selected, and their kernels were used for the phenotyping of the kernel width.

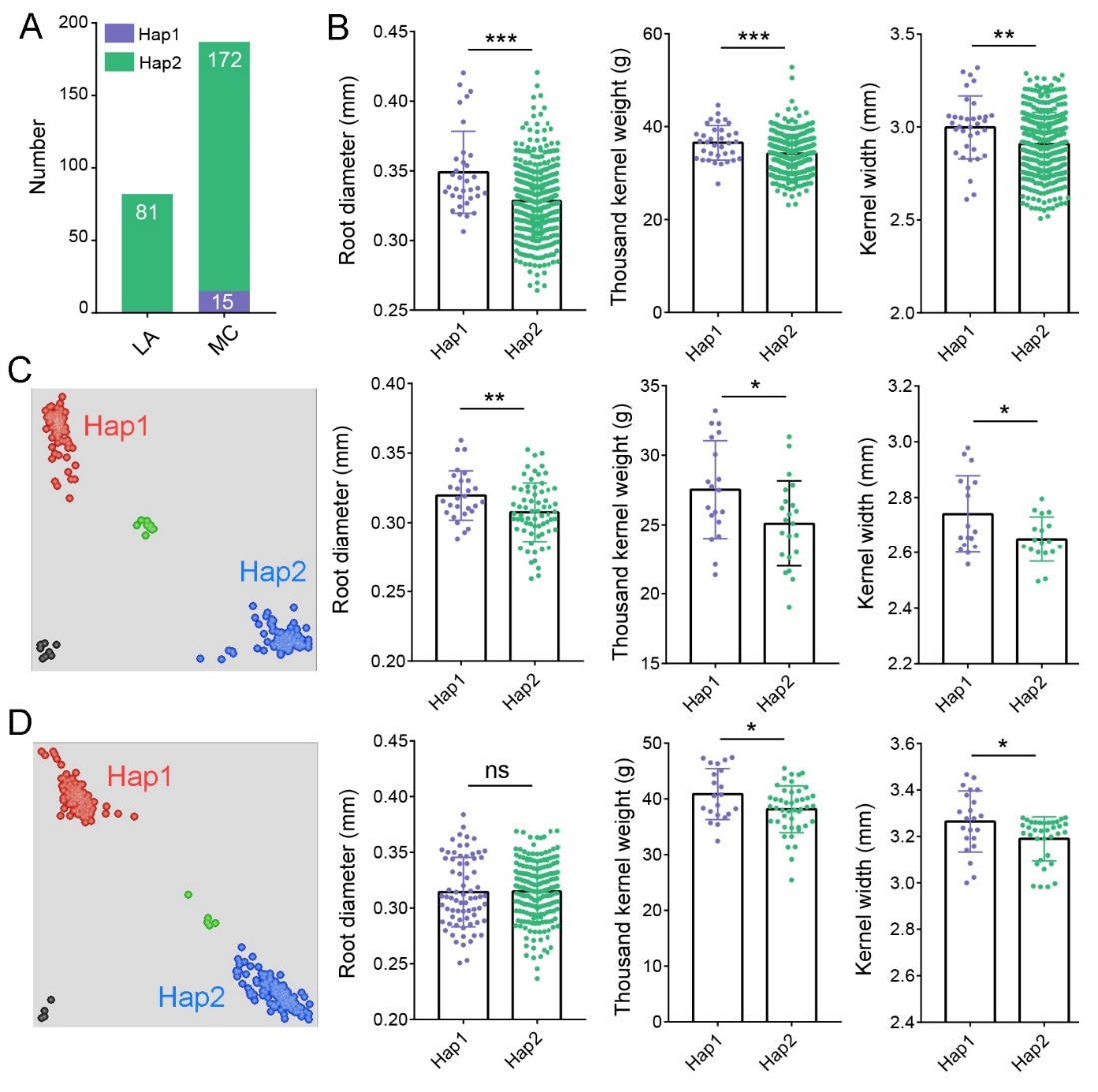

**Supplemental Figure 4. Validation of the genetic effects of bi-effect block b047563 on root diameter, thousand kernel weight and kernel width.** (A) Frequency of Hap1 and Hap2 in LA and MC groups. (B) The phenotypic value distributions and significance values (Student's *t*-tests) between genotypes carrying Hap1 and Hap2 of block b047563 in the natural association population (top panels); F_5:6_ RIL populations derived from crosses between MK95 (Hap 1) and Chinese Spring (Hap2) (central panels) and between MK147 (Hap 1) and Chinese Spring (Hap2) (lower panels). Asterisks indicate significance: * < 0.05; ** < 0.01; *** < 0.001. (C-D) The KASP assays to distinguish Hap1 and Hap2 genotypes in F_5:6_ RIL populations derived from crosses between (C) MK95 (Hap 1) and Chinese Spring (Hap2) and (D) between MK147 (Hap 1) and Chinese Spring (Hap2). For each F_5:6_ RIL population, five to eight plants of each line were genotyped, and the homozygous plants were phenotyped for the root-related traits. The homozygous lines were selected, and their kernels were used for the phenotyping the thousand kernel weight and kernel width.

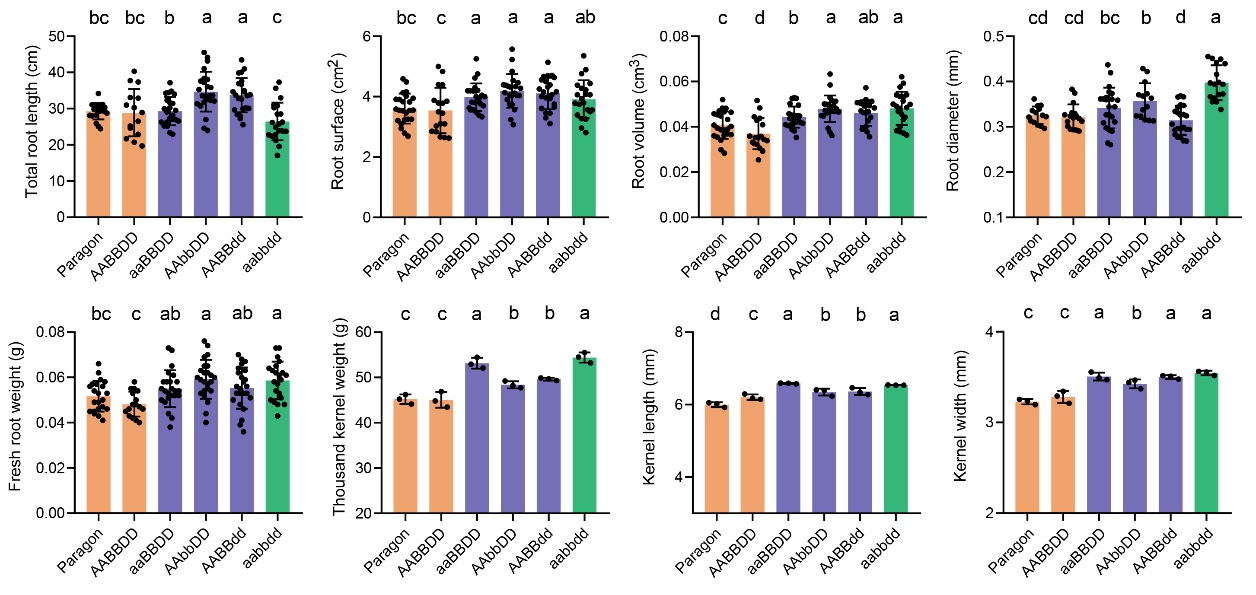

**Supplemental Figure 5. The root and kernel phenotypes of wild type (Paragon and AABBDD), *TaGW2* single mutants (aaBBDD, AAbbDD, and AABBdd) and triple mutant (aabbdd).** Data are means ± SD; letters on top indicate statistical differences (*P-*value < 0.05) based on the least significant difference (LSD) test. Root phenotypes were evaluated at 14 days after germination.

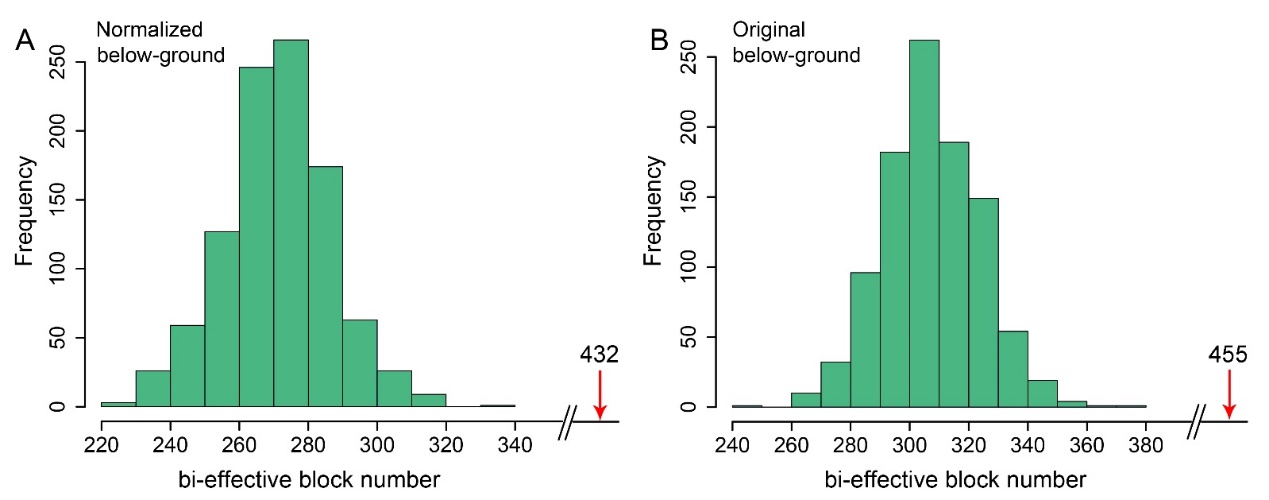

**Supplemental Figure 6. The distribution of bi-effect block numbers in the 1,000 permutation tests.** For each test, 3,389 blocks (equal to the number of selection blocks) were randomly sampled from genome-wide blocks and the bi-effect block number was recorded. Both the normalized values of below-ground traits (A) and original values of below-ground traits (B) were used for the permutation tests, and the bi-effect block numbers derived from normalized and original below-ground traits were both smaller than the observed numbers (red arrows).

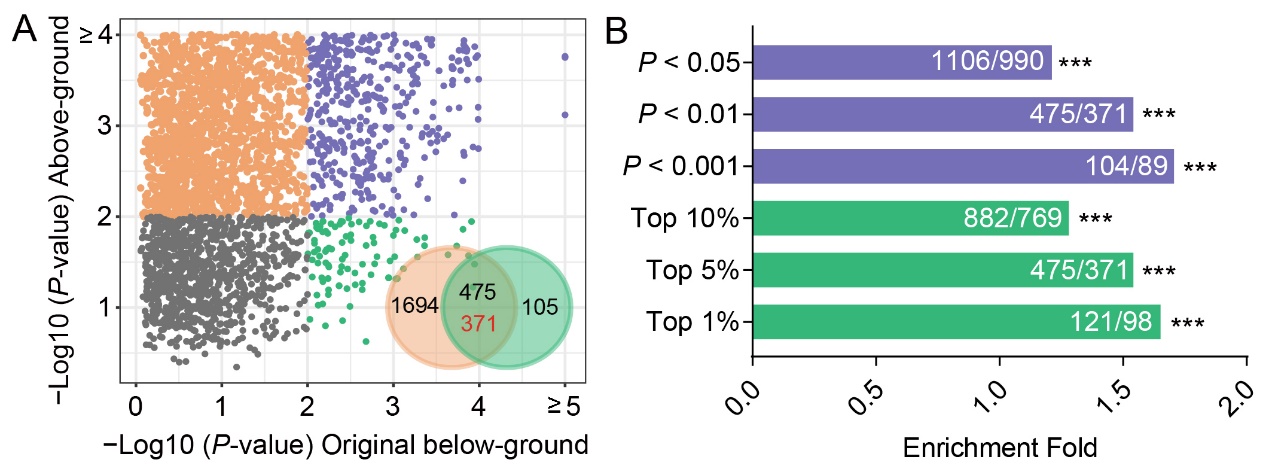
 **Supplemental Figure 7. The original phenotype values of below-ground traits (without adjusting for kernel weight) were used for the enrichment analysis.** (A) Each point represents a selection block. The *x*- and *y*-axis indicate the *P-*value of each selection block in the genetic effect estimations. In the Venn diagrams, the orange and green sections represent the selection blocks which had genetic effects on the above- and below-ground traits, respectively. The overlap represents the bi-effect blocks. The numbers in black are the observed values, while in red are the maximum of the expected values in the 1,000 permutation tests. (B) The fold-enrichment of bi-effect blocks in the selection sweeps increased with more stringent thresholds, from *P-*value 0.05 to 0.001 in the genetic effect estimation and from top 10% to 1% in the selection sweep detections. For the number pair on each bar, the left number is the observed value, and the right number is the maximum of the expected values in the 1,000 permutation tests.

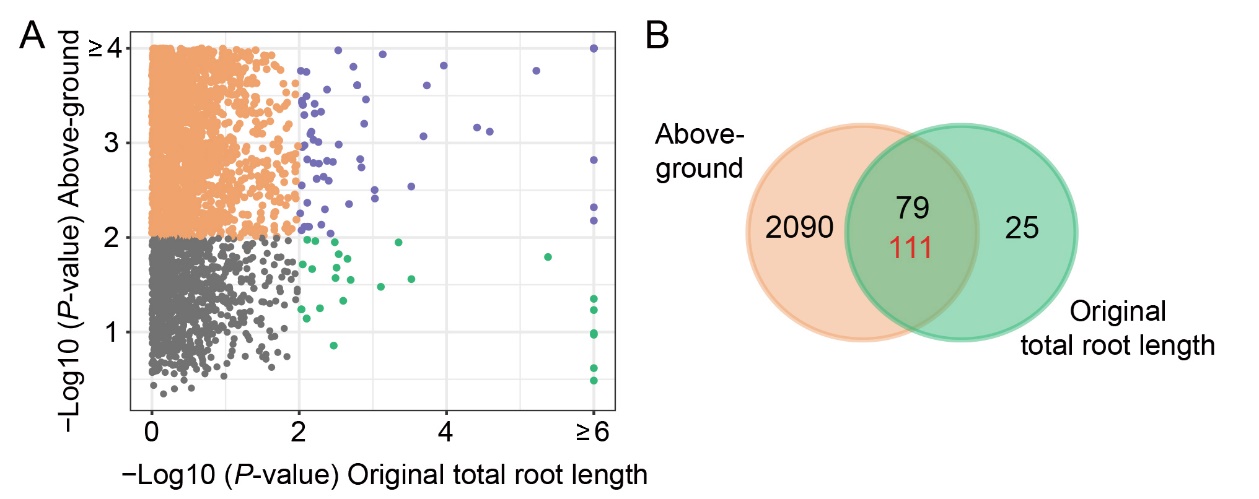

**Supplemental Figure 8.** **The bi-effect blocks that have genetic effects on the above-ground traits and original values of total root length (without adjusting for kernel weight) were not enriched in the selection blocks.** (A) The *x*- and *y*-axis indicate the *P-*value of each selection block in the genetic effect estimations. (B) The numbers of bi-effect blocks that have genetic effects on the above-ground traits and original total root length were plotted as orange and green, respectively. The observed value of the bi-effect block is 79, and the maximum of expected values is 111 in the 1,000 permutation tests.

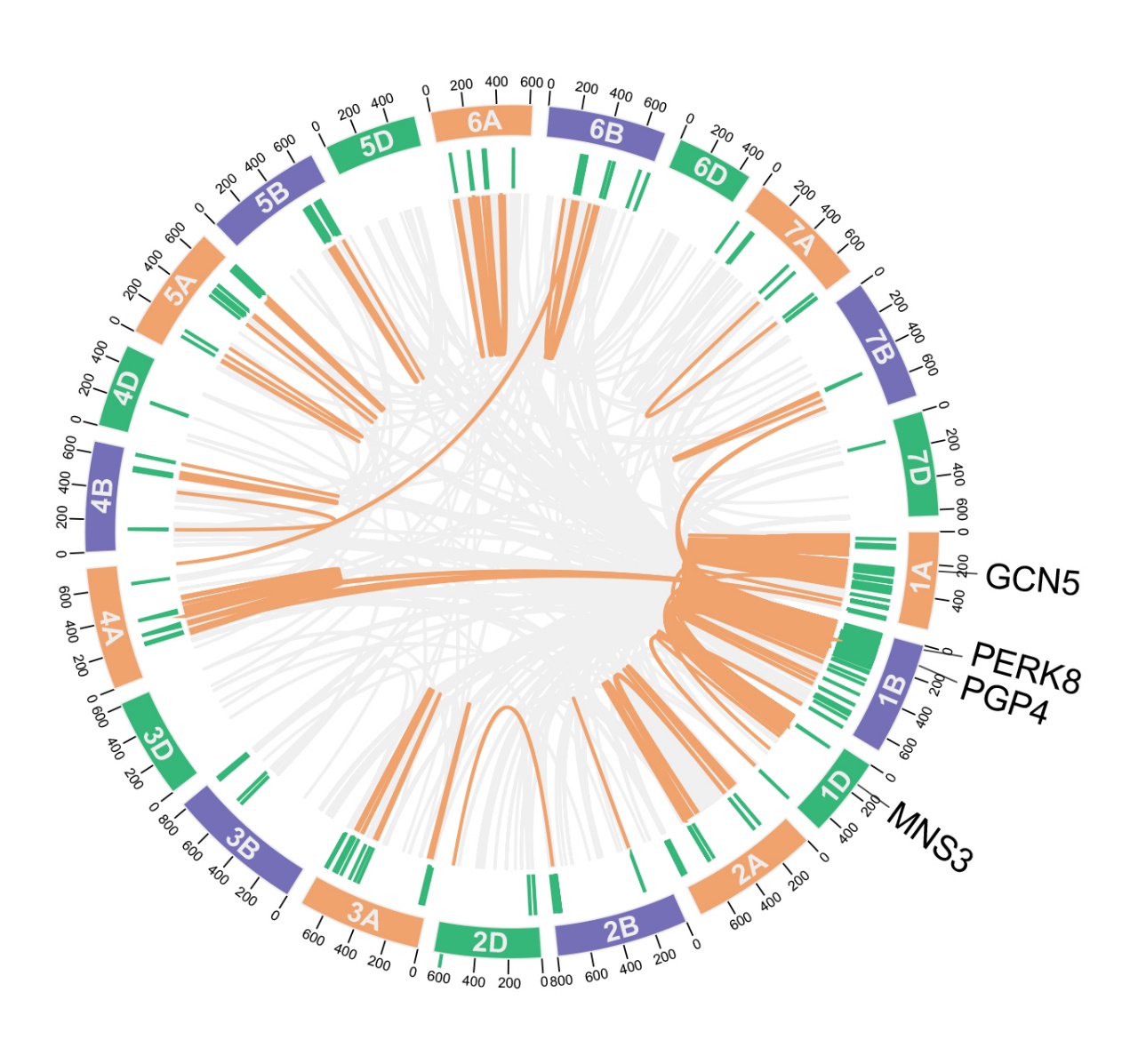

**Supplemental Figure 9. The bi-effect selection blocks regulate the expressions of root development-related genes with eQTLs.** Outer circle depicts 21 wheat chromosomes with Mbp physical position. The green lines in the second layer represent the physical positions of bi-effect selection blocks. The orange and grey curves in the center represent relationships between eQTLs and their regulating genes, in which the orange curves indicate that the regulated genes were significantly associated with below-ground trait variations in the TWAS.

**Supplemental Data**

Supplemental Data 1. Phenotype values of all accessions used in this study.

Supplemental Data 2. Genetic effects of genomic linkage blocks on the investigated traits.

Supplemental Data 3. Selection sweeps detected between LA and MC groups.

Supplemental Data 4. Well-known selected genes in the LD blocks.

Supplemental Data 5. Genetic effects of the selected haplotypes (with higher frequency in the MC group) in the bi-effect selection blocks on above- and below-ground traits.

Supplemental Data 6. The eQTLs located in bi-effect selection blocks.

Supplemental Data 7. The 276 genes whose expression levels was regulated by the bi-effect selection blocks with eQTLs and were significantly associated with below-ground traits in TWAS.

Supplemental Data 8. The KASP primer sequence.
